# Taxonomic and functional shifts in the perinatal gut microbiome of rhesus macaques

**DOI:** 10.1101/2022.03.02.482686

**Authors:** Nicholas S. Rhoades, Isaac R. Cinco, Sara M. Hendrickson, Mark K. Slifka, Ilhem Messaoudi

## Abstract

Pregnancy and the postpartum period result in some of the most dramatic metabolic, hormonal, and physiological changes that can be experienced by an otherwise healthy adult. The timing and magnitude of these changes is key for both maternal and fetal health. One of the factors believed to critically modulate these physiological changes is the maternal gut microbiome. However, the dynamic changes in this community during the perinatal period remain understudied. Clinical studies can be complicated by confounding variables such as diet and other drivers of heterogeneity in the human microbiome. Therefore, in this study, we conducted a longitudinal analysis of the fecal microbiome obtained during the perinatal and post-partum periods in 25 rhesus macaques using 16S rRNA gene amplicon sequencing and shotgun metagenomics. Shifts at both the taxonomic and functional potential level were detected when comparing pregnancy to postpartum samples. Taxonomically, *Alloprevotella, Actinobacillus,* and *Anaerovibrio* were enriched during pregnancy while *Treponema,* Lachnospiraceae, and *Methanosphaera* were more abundant post-partum. Functionally, pregnancy was associated with increased abundance in the pathway to produce the beneficial short chain fatty acid (SCFA), butyrate, while pathways associated with starch degradation and folate transformation were more abundant postpartum. These data demonstrate dramatic changes in the maternal gut even in the absence of dietary changes and suggest that rhesus macaques could provide a valuable model to determine how changes in the microbiome correlate to other physiological changes in pregnancy.

## INTRODUCTION

The perinatal period, which encompasses pregnancy up to 1-year postpartum, is characterized by large physiological, immunological, and hormonal changes that impact maternal and fetal health. This shift is characterized by an increase in insulin resistance, along with increased leptin and adiponectin (1, 2) that are critical to ensure that the fetus receives adequate nutrition and preparing the mother for metabolic demands imposed by lactation. A disruption of these metabolic adaptations can lead to adverse outcomes such as gestational diabetes and pre-eclampsia (3–5), which in turn can have a negative impact on the infant including high incidence of preterm birth, increased incidence of infection, large for gestational age (LGA) and reduced cognitive development (6–8). One of the factors that significantly modulate maternal metabolism but remains understudied during the perinatal period is the gut microbiome (9).

The gut is the most densely populated microbial community in the human body that is composed of bacterial symbionts, as well as archaeal, fungal and viral members (10). The gut microbiome plays critical roles in vitamin production (11), immune homeostasis (12), and metabolism of indigestible substrates among others (13). This community stabilizes in adulthood (14), but sensitive to environmental factors such as dietary shifts (15), antibiotic use (16), and other shifts in host physiology such as pregnancy (18). Early studies found that the pregnancy gut microbiome is less diverse, more variable, and harbors an increased abundance of Proteobacteria (19). However, more recent studies found that the microbiome remained stable during pregnancy, but shifted significantly in the postpartum period (20). These shifts are believed to be the result of the maternal metabolic and hormonal changes required for a successful pregnancy. Additionally, stress during pregnancy can exacerbate this dysbiotic microbiome state (21), which has also been implicated in postpartum depression (22).

Another key reason for understanding the perinatal maternal gut microbiome is the vital role it plays in seeding the infant microbiome (23). The infant is first exposed to maternal vaginal and fecal microbes at birth. The development and maintenance of the infant gut microbiome has long-term ramification for maturation of the immune system (24), protection from enteric infection (25), and establishment of a healthy metabolic state (26). The establishment of the infant microbiome is shaped by influenced by a multitude of factors such as delivery method (28), antibiotic use (29), and breastfeeding (30).

Rhesus macaques are a valuable preclinical model to study the role of the microbiome in health and disease as they have a gut microbiome similar to humans, especially those in the developing world (31, 32). Moreover, rhesus macaques are a vital model for the study of perinatal and reproductive health (33, 34). In this study, we utilize a combination of 16S rRNA amplicon, and shotgun metagenomic sequencing to longitudinally characterize the fecal microbiome of rhesus macaques during the perinatal period. We observed both taxonomic and functional potential shifts within the fecal microbiome both within individual animals and across our entire study population before and after delivery. Taxonomically, *Alloprevotella, Actinobacillus,* and *Anaerovibrio* were enriched during pregnancy while *Treponema,* Lachnospiraceae, and *Methanosphaera* were more abundant post-partum. Shift in the functional potential of the perinatal gut microbiome included an increased abundance in the pathway to produce the beneficial short chain fatty acid (SCFA), butyrate, during pregnancy, and pathways associated with starch degradation and folate transformation postpartum.

## RESULTS

### Taxonomic shifts in the perinatal gut microbiome

We utilized 16S rRNA gene amplicon sequencing of rectal swabs collected prebirth (~90 and 60 days prior to birth) and post-birth (~30 and 90 days after giving birth) (see experimental design **Figure 1A**) to determine shifts in microbial communities between pregnancy and postpartum. Prior to analysis we confirmed that samples were free of contamination PCR bias using negative controls and sequenced community standards (**Figure S1A**). All samples were dominated by Bacteroidetes *(Prevotella,* Rikenellaceae*, Alloprevotella)* and Firmicutes (Lachnospiraceae, *Lactobacillus, Streptococcus*) along with Proteobacteria (*Helicobacter and Campylobacter*) and Spirochetes *(Treponema)* (**Figure 1B**). Despite the lack of changes in housing or diet, we observed a distinct shift in the overall composition of the maternal gut microbiome when comparing samples collected during pregnancy and postpartum (**Figure 1C**). At both timepoints measured during pregnancy, the maternal gut microbiome was more variable than postpartum (**Figure 1D**). Additionally, samples collected postpartum had an increased number of observed amplicon sequencing variants (ASVs) when compared to samples collected during pregnancy (**Figure 1E)**. This pattern was observed across the entire cohort and when we conducted a pairwise comparison for animals that had samples from all 4 time points (**Figure S1B**).

**Figure 1:**
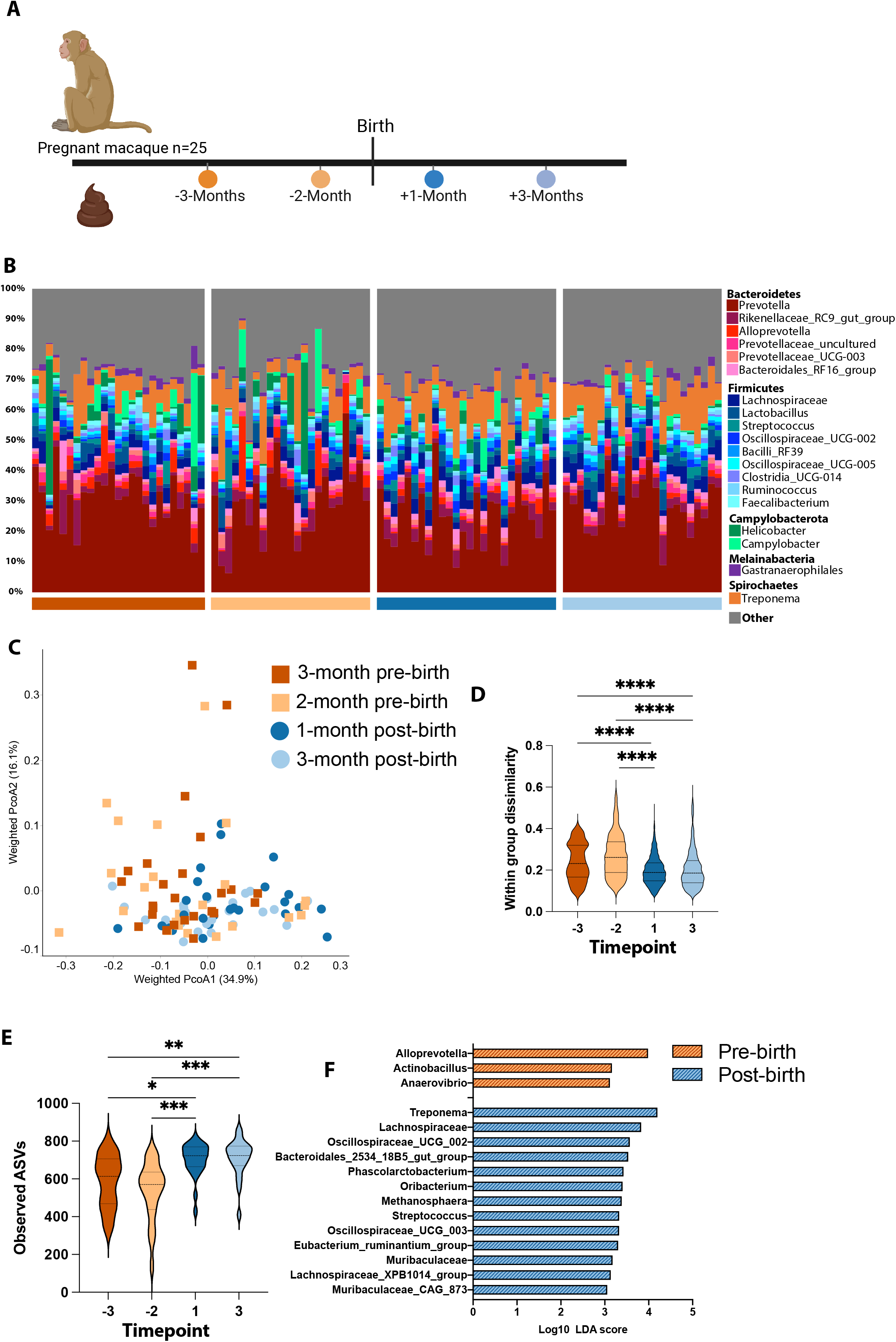
Landscape and perinatal shifts of the maternal gut microbiome. (A) Study design. (B) Stacked bar plot organized by time-point. All taxa below 1% average abundance grouped into the “Other” category. Each vertical bar represents a single sample. (C) Principal coordinate analysis (PcoA) of weighted UniFrac distances between microbial communities colored by timepoint. (D) Violin plot of weighted UniFrac distances between the fecal microbiome sample collected at the same timepoint. (E)Violin plot of observed amplicon sequencing variants at each time point. (F) Differentially abundant taxa between pregnant and post-partum samples with pregnancy vs. post-partum used as the class and individual time-points as sub-class. Differential abundance was determined using LEFsE (Log_10_ LDA score > 2). Significance for panels B & D was determined using 1-way ANOVA with post-hoc Sidak multiple comparisons test ** p < 0.01, *** p < 0.001, **** p < 0.0001. Solid lines within each violin indicate the median value along with the 25^th^, and 75^th^ percentiles for that timepoint.

We next compared pre- and postpartum samples using linear discriminant analysis effect size (LEfSe) to determine which taxa were driving the observed differences in gut microbiome composition. During pregnancy, the maternal gut microbiome was enriched in *Alloprevotella, Actinobacillus,* and *Anaerovibrio* (**Figure 1F**), while the postpartum gut microbiome was enriched in *Treponema*, multiple Lachnospiraceae, and *Methanosphaera* among others (**Figure 1F**). The enrichment of *Alloprevotella* in pregnancy samples was driven by a significantly higher abundance in early pregnancy samples when compared to postpartum samples (**Figure S1C, D**). On the other hand, relative abundance of *Treponema* was lower during pregnancy (**Figure S1E, F**). While the shift in these higher abundance genera were significant, we observed more dramatic shifts in the abundance of lower abundant taxa postpartum. For example, Oscillospiraceae_UCG002 was transiently more abundant 1-month postpartum, while *Methanosphaera* was increased at both postpartum timepoints (**Figure S1G-J**).

### Shotgun metagenomics confirm taxonomic trends and reveals perinatal shift in the perinatal gut microbiome

To further explore how these taxonomic shifts impacted the metabolic potential of the gut microbiome, we utilized shotgun metagenomics. Shotgun metagenomic libraries were prepared from a subset of fecal samples collected 2-months prior to birth and 1-month postpartum (n=15/timepoint). Two postpartum libraries had less than 1 million reads after host decontamination and were excluded from future analysis. In contrast to the 16S rRNA gene amplicon sequencing data, the overall taxonomic composition assessed by shotgun metagenomics did not significantly differ between the timepoints (**Figure 2A**). This disagreement could be due to lower sample numbers used for our shotgun metagenomic experiment or potentially driven by the lack of phylogenetically informed beta diversity metrics (UniFrac) for shotgun metagenomic data. Nevertheless, we were able to identify multiple differentially abundant species between these two timepoints. For example, abundance of *Prevotella* species, the most abundant genera from our 16S data, shifted from an enrichment of *Prevotella copri* and *Prevotella* sp_AM42_24 pre-birth to *Prevotella* sp_CAG_873 postpartum (**Figure 2B**). Species that were enriched in postpartum samples included *Oscillibacter* sp_57_20, *Phasolarctobacterium succinatutens* and *Treponema succinifaciens* (**Figure 2B**).

**Figure 2:**
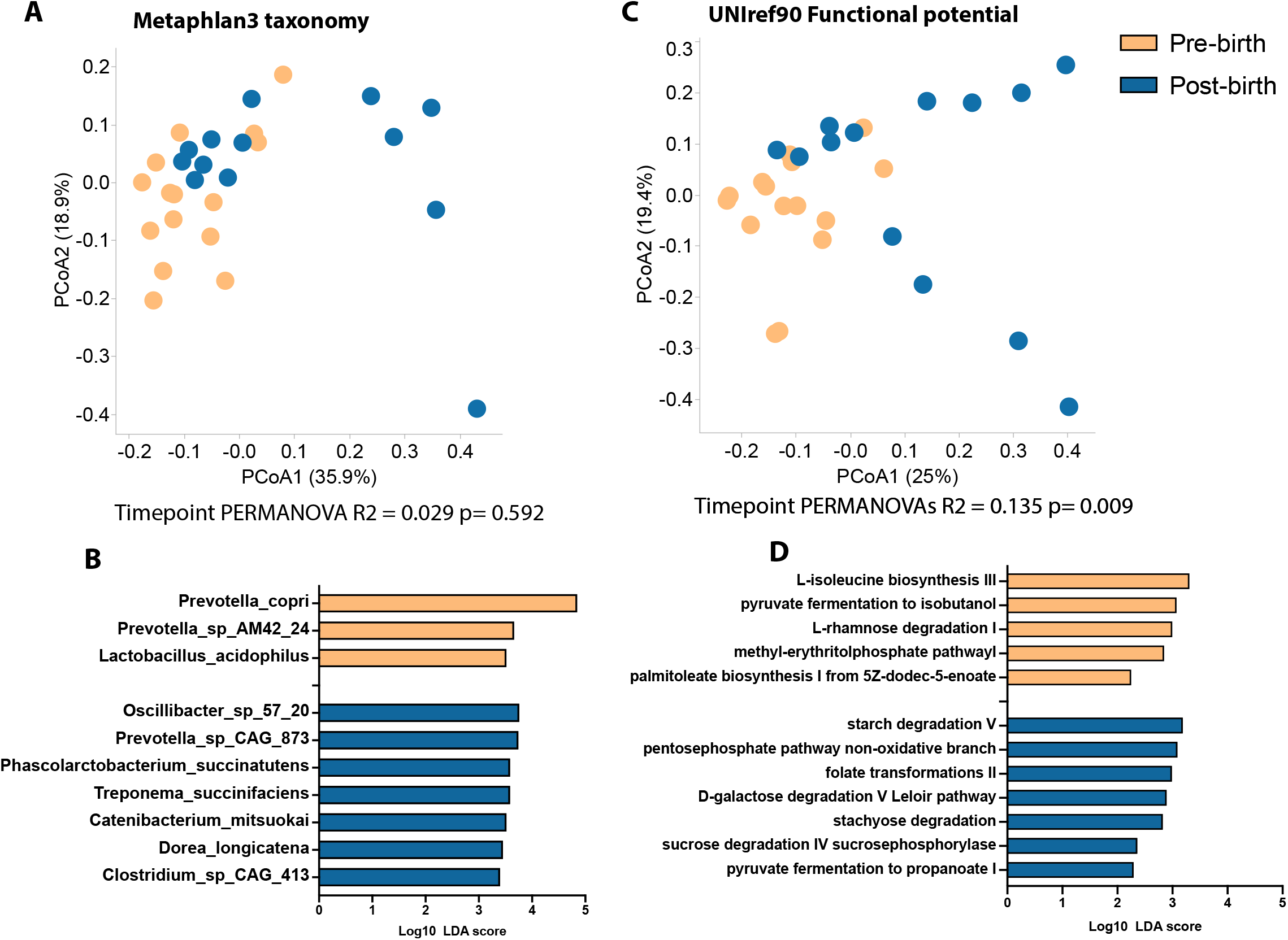
Perinatal shifts in the functional potential and species level taxonomy of the maternal microbiome. (A) PcoA of Bray-Curtis dissimilarity built on the abundance of all functional genes annotated using humann3 and the Uniref90 database. Colored by timepoint. (B) Differentially abundant Metacyc pathways between pre- and post-birth samples (LEfSe, Log_10_ LDA score > 2). (C) Principal coordinate analysis (PcoA) of Bray-Curtis dissimilarity built on species-level abundance from MetaPhlan3. Colored by timepoint. (D) Differentially abundant species between pre- and post-birth samples (LEfSe, Log_10_ LDA score > 2).

We also used shotgun metagenomics to assess the functional potential of the pregnant rhesus gut microbiome (**Figure 2C**). In total, 77 Metacyc pathways were differentially abundant between the two timepoints. More specifically, the functional capacity of the late pregnancy microbiome showed enrichment of “pyruvate fermentation to isobutanol”, “methyl-erythritol phosphate pathway”, and “L-isoleucine biosynthesis” pathways (**Figure 2D**). In contrast, the early postpartum gut microbiome had a higher abundance of “Starch degradation”, “folate transformation”, and “pyruvate fermentation to propionate” (**Figure 2D**).

To better understand which microbial functions best distinguished the pre- and post-delivery time points, we used a supervised random forest model built using gene ontology (GO) terms. The overall random forest model was 82% accurate at classifying samples into the two timepoints (**Figure 3A**). Next, we extracted and plotted the abundance of the 25 GO terms that best distinguished the two timepoints (**Figure 3B**).

**Figure 3:**
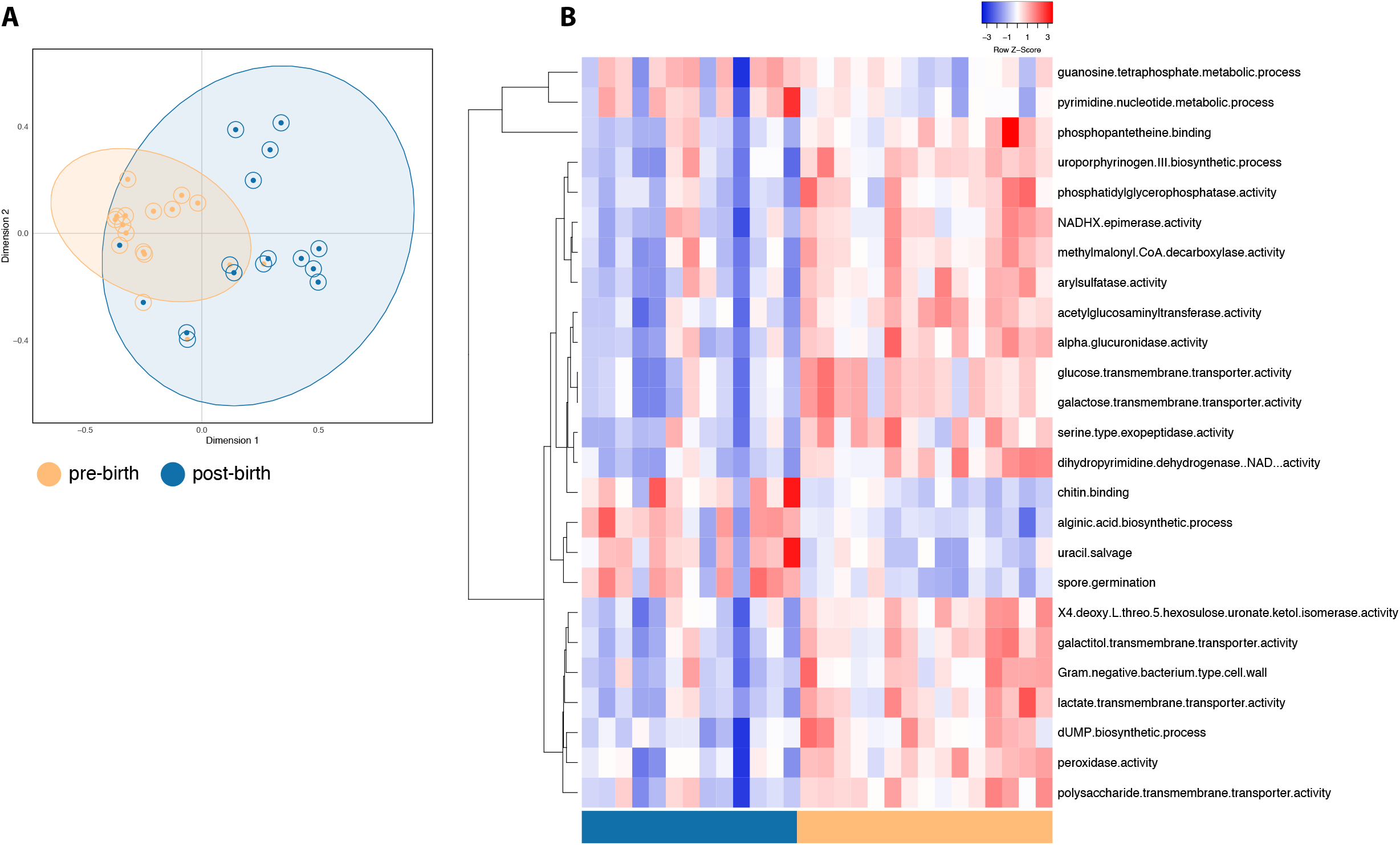
Random Forest highlights shifts in perinatal gut microbiome functional potential. (A) MDS plot of proximity scores from a random forest of gene ontology (GO) terms colored by timepoint. Inner circles denote samples actual group. Outer circles denote if a sample was misclassified at any point during the random forest model generation. The color of the outer circle represents which group that sample was misclassified as. (B) Heat map of the 25 most important GO terms as predicted by random forest modeling.

Many of the GO terms that were more abundant in pre-birth samples were associated with metabolism and transport of carbohydrates notably “alpha glucuronidase activity”, “galactose transmembrane transporter activity”, and “lactate transmembrane transporter activity” (**Figure 3B**). Interestingly some of the GO terms that best distinguish postpartum samples are associated with fungi rather than bacteria such as “spore germination” and “chitin binding” (**Figure 3B**).

## DISCUSSION

Pregnancy and the perinatal period encompass some of the most dramatic shifts in metabolism, immunity, and hormonal balance that a healthy adult can experience. It is well established that the gut microbiome can modulate these shifts. Indeed, a recent study of a cohort in rural Zimbabwe found that the taxonomic composition of the maternal fecal microbiome was a strong predictor of gestational age, birth weight, and neonatal growth (46). However, our understanding of the changes that the maternal gut microbiome undergoes during the perinatal period is limited. In this study, we explored taxonomic and functional shifts in the gut microbiome of rhesus macaques during the perinatal period. One limitation of this study that pre-pregnancy and early pregnancy samples (1^st^ trimester) were not collected, which precluded us from fully capturing the dynamics of the changes in the maternal microbiome.

Our data indicate that the maternal microbiome was less rich (lower observed ASVs) but more heterogeneous (increased within group dissimilarity, beta diversity) during pregnancy. This increase in beta-diversity with pregnancy is in line with data from multiple human and animal studies, particularly late in pregnancy (18–20, 37). On the other hand, previous clinical studies have reported a drop, an increase, or no change in the alpha diversity of the maternal microbiome (18, 19, 35) likely due to the high heterogeneity within the healthy human microbiome (36). Some of the previously reported changes include increased relative abundance of Actinobacteria, Firmicutes, and Proteobacteria during late and post pregnancy, while that Bacteroidetes decreased (19, 20). We observed no significant changes in the abundance of taxa belonging to Actinobacteria or Proteobacteria. However, we did observe an increase in multiple Firmicutes post-birth including Lachnospiraceae, *Phasolarctobacterium*,

Oscillospiraceae and *Eubacterium* among others. Many of these taxa are associated with the production of beneficial SCFA within the gut (38, 39), which is linked to lower blood pressure and reduced preeclampsia risk (47, 48). We also observed perinatal shifts in the abundance of genera *Treponema*, *Methanosphaera*, and *Alloprevotella*. These taxa are uncommon in the human gut microbiome in the developed world but common in humans from low- and middle-income countries, especially humans living in a rural rather than urban setting (40, 41) further supporting the notion that gut microbiome of rhesus macaques is more similar to that of humans in the developing world (31,42).

Many of the perinatal taxonomic trends we observed at both the 16S amplicon and shotgun metagenomic level were driven by taxa with unique metabolic capacity such as methanogenic *Methanosphaera,* and fiber-degrading *Treponema,* thus suggesting that the metabolic capacity of the prenatal gut microbiome changes during the perinatal period. Many of our findings including the increased potential to produce Butyrate in late pregnancy agree with what has been observed in humans (49). We also observed increased enrichment of GO terms associated with the transportation of simple sugars such as Lactate, glucose and galactose in late pregnancy compared to post birth samples. Postpartum, the gut microbiome was enriched in pathways such as starch degradation, folate transformation, and stachyose degradation that are important for the degradation of complex substrates and production of secondary metabolites rather than the metabolism of simple sugars observed in late pregnancy. This switch is likely a reflection of the host nutritional needs. Additionally, our random forest analysis revealed that the post-birth gut microbiome was enriched in GO-terms associated with fungal colonization of the gut including spore germination, and chitin binding. To our knowledge, the gut mycobiome has not previously been examined during the perinatal period.

## METHODS

### Cohort description

Longitudinal samples were obtained from a total of 25 reproductive age rhesus macaques at 4 timepoints, 17 of which had sample collected at every timepoint. These timepoints included including: 3-months pre-birth (90.2 +/− 20.5 days pre-birth), 2-months pre-birth (60.0 +/− 20.0 days pre-birth), 1-month post-birth (29.8 +/− 1.5 days post-birth), and 3-months post-birth (91.2 +/− 4.0 days post-birth). All rhesus macaque studies were overseen and approved by the OHSU/ONPRC Institutional Animal Care and Use Committees (IACUC) per the National Institutes of Health guide for the care and use of laboratory animals. Animals were housed per the standards established by the US Federal Animal Welfare Act and *The Guide for the Care and Use of Laboratory Animals*. All animals were tested for simian viruses (Simian Immunodeficiency Virus, Simian Retrovirus 2, Macacine herpesvirus 1, and Simian T lymphotropic virus) and received a tuberculin test semi-annually. All animals were vaccinated against *Campylobacter coli* at both pre-birth time points for an unrelated study.

### 16S amplicon sequencing

Total DNA was extracted from rectal swabs using the DNeasy Powersoil Pro Kit (Qiagen, Valencia, CA, USA). The hypervariable V4-V5 region of the 16S rRNA gene was amplified using PCR primers (515F/926R with the forward primers including a 12- bp barcode). PCR reactions were conducted in duplicate and contained 12.5 μl GoTaq master mix, 9.5 μl nuclease-free H_2_0, 1 μl template DNA, and 1 μl 10 μM primer mix. Thermal cycling parameters were 94°C for 5 minutes, 35 cycles of 94°C for 20 seconds, 50°C for 20 seconds, 72°C for 30 seconds, followed by 72°C for 5 minutes. PCR products were purified using a MinElute 96 UF PCR Purification Kit (Qiagen, Valencia, CA, USA). Libraries were sequenced (2 x 300 bases) using an Illumina MiSeq.

Raw FASTQ 16S rRNA gene amplicon sequences were uploaded and processed using the QIIME2 analysis pipeline (50). Briefly, sequences were demultiplexed and the quality filtered using DADA2 (51), which filters chimeric sequences and generates an amplicon sequence variant (ASV) table equivalent to an operational taxonomic unit (OTU) table at 100% sequence similarity. Sequence variants were then aligned using MAFFT (52) and a phylogenetic tree was constructed using FastTree2 (53). Taxonomy was assigned to sequence variants using q2-feature-classifier against the SILVA database (release 138) (54). To prevent sequencing depth bias samples were rarified to 13,781 sequences per sample before alpha and beta diversity analysis. QIIME 2 was also used to generate the following alpha diversity metrics: richness (as observed ASV), Shannon evenness, and phylogenetic diversity. Beta diversity was estimated in QIIME 2 using weighted and unweighted UniFrac distances (55).

### Shotgun metagenomic library preparation and analysis

Shotgun metagenomic libraries were prepared from 100 ng of gDNA using the iGenomx RIPTIDE (iGenomx, South San Fransisco CA) per iGenomx recommended protocol and sequenced on an Illumina HiSeq 4000 2■×■100. Raw demultiplexed reads were quality filtered using Trimmomatic (56), and potential host reads were removed by aligning trimmed reads to the *Macaca mulata* genome (Mmul 8.0.1) using BowTie2 (57). Trimmed and decontaminated reads were then annotated using the HUMAnN3 pipeline using default settings with the UniRef90 database and assigned to Metacyc pathways. Functional annotations were normalized using copies per million (CPM) reads before statistical analysis (58–60). Species-level taxonomy was assigned to quality-controlled short reads using Metaphlan3 (61).

### Statistics

PERMANOVAs were performed using the Vegan (62) function ADONIS. 1-way, non-parametric Kruskal-Wallis ANOVA were implemented using PRISM (V8) to generate p-values and utilizing the Dunns post-hoc-test when the initial ANOVA was significant. The LEfSe algorithm was used to identify differentially abundant taxa and pathways between groups with a logarithmic Linear discriminant analysis (LDA) score cutoff of 2 (63)

## ACKNOWLEDGMENTS

We would like to thank the ONPRC animal husbandry team for their assistance with sample collections and animal care throughout this study.

---

**Figure S1:**
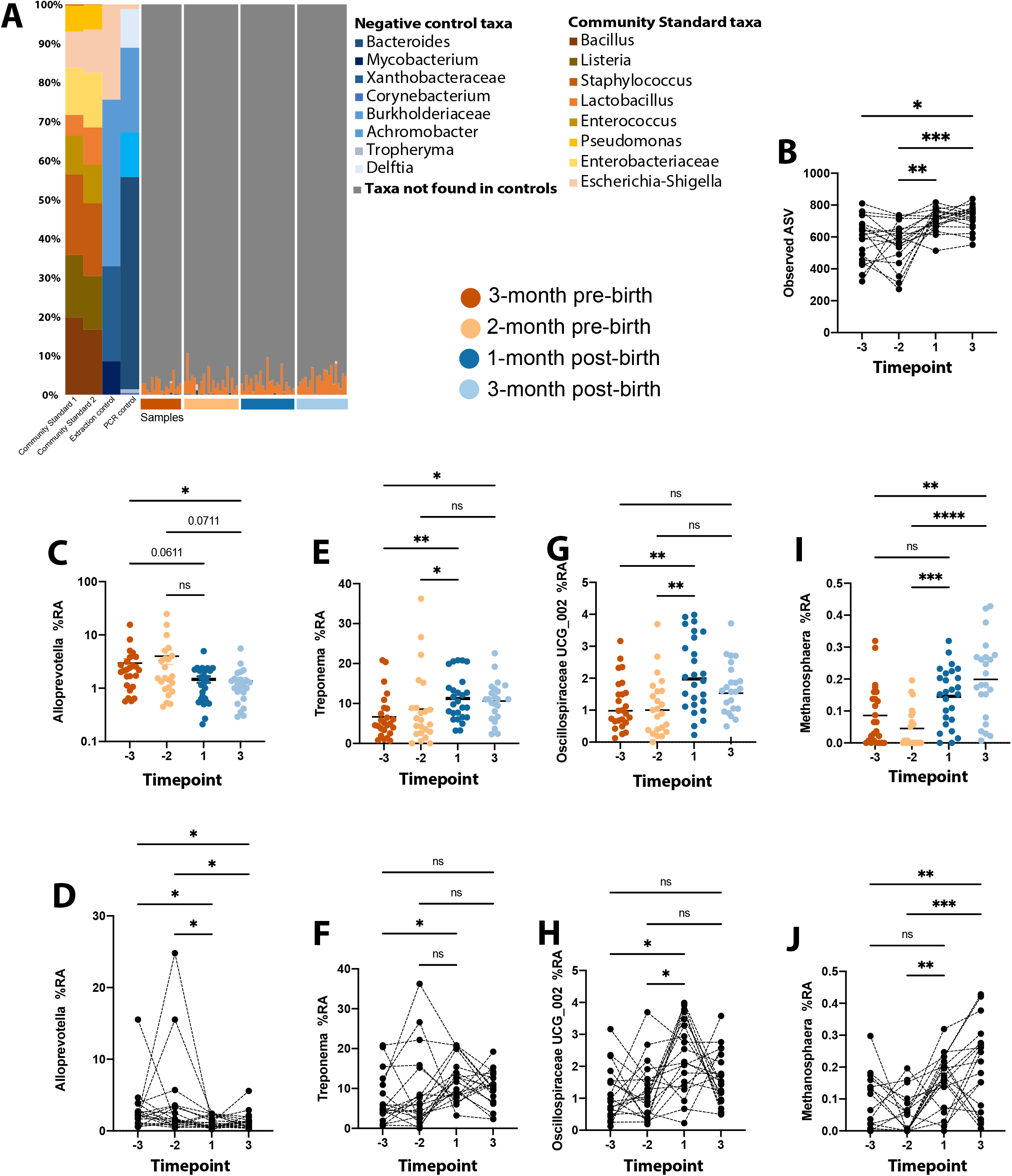
Additional taxonomic comparisons and pairwise comparisons. (A) Stacked bar plot comparing microbial communities of control samples to the biological samples included in this study. (B-J) Scatterplot of (B) observed ASV (C, D) Alloprevotella, (E, F) Treponema, (G, H) Oscillospiraceae_UCG002 and (I, J) Methanosphaera. Significance for B, D, F, H and J was measured by nonparametric one-way repeated-measure ANOVA (Friedman test) with Dunn’s *post hoc* comparisons between time points. Bars denote significance of *post hoc* tests: *, *P* < 0.05; **, *P* < 0.01; and ***, *P* <0.001. Each dot represents an individual sample, with solid lines connecting samples from the same individual across time. Significance for panels C, E, G, I was determined using 1-way ANOVA with post-hoc Sidak multiple comparisons test ** p < 0.01, *** p < 0.001, **** p < 0.0001. Each dot represents an individual sample colored by timepoint.

